# Phylogenomic analysis of the Neocallimastigomycota: Proposal of *Caecomycetaceae* fam. nov., *Piromycetaceae* fam. nov., and emended description of the families *Neocallimastigaceae and Anaeromycetaceae*

**DOI:** 10.1101/2022.07.04.498725

**Authors:** Radwa A. Hanafy, Yan Wang, Jason E. Stajich, Carrie J. Pratt, Noha H. Youssef, Mostafa H. Elshahed

## Abstract

The anaerobic gut fungi (AGF) represent a coherent phylogenetic clade within the Mycota. Twenty genera have been described so far. Currently, the phylogenetic and evolutionary relationships between AGF genera remain poorly understood. Here, we utilized 53 transcriptomic datasets from 14 genera to resolve AGF inter-genus relationships using phylogenomics, and to provide a quantitative estimate (amino acid identity) for intermediate rank assignments. We identify four distinct supra-genus clades, encompassing genera producing polyflagellated zoospores, bulbous rhizoids, the broadly circumscribed genus *Piromyces*, and the *Anaeromyces* and affiliated genera. We also identify the genus *Khoyollomyces* as the earliest evolving AGF genus. Concordance between phylogenomic outputs and RPB1 and D/D2 LSU, but not RPB2, MCM7, or ITS1, phylogenies was observed. We combine phylogenomic analysis, and AAI outputs with informative phenotypic traits to propose accommodating 13/20 AGF genera into four families: *Caecomycetaceae* fam. nov. (encompassing genera *Caecomyces* and *Cyllamyces*), *Piromycetaceae* fam. nov. (encompassing the genus *Piromyces*), emend the description of fam. *Neocallimastigaceae* to only encompass genera *Neocallimastix*, *Orpinomyces*, *Pecramyces*, *Feramyces*, *Ghazallomyces*, and *Aestipascuomyces*, as well as the family *Anaeromycetaceae* to include the genera *Oontomyces*, *Liebetanzomyces*, and *Capellomyces* in addition to *Anaeromyces*. We refrain from proposing families for the deeply branching genus *Khoyollomyces*, and for genera with uncertain position (*Buwchfawromyces, Joblinomyces*, *Tahromyces, Agriosomyces, Aklioshbomyces,* and *Paucimyces*) pending availability of additional isolates and sequence data. Our results establish an evolutionary- grounded Linnaean taxonomic framework for the AGF, provide quantitative estimates for rank assignments, and demonstrate the utility of RPB1 as additional informative marker in Neocallimastigomycota taxonomy.

## Introduction

Members of the anaerobic gut fungi (AGF) represent a phylogenetically, metabolically, and ecologically coherent clade in the kingdom Mycota [1]. Twenty genera and thirty-six different species have been described so far [2]. A recent review has provided detailed description of current genera and resolved historical inaccuracies and synonymies within the *Neocallimastigomycota* [2]. Further, criteria for the identification and characterization, as well as guidelines for genus- and species-level rank assignment for novel AGF isolates have recently been formulated [3]. In spite of such progress, the phylogenetic and evolutionary relationships between various genera within the Neocallimastigomycota are currently unclear. Similarities in specific microscopic traits (zoospore flagellation, thallus development, and rhizoidal growth patterns) across genera have been identified; and the significance of using such traits for proposing higher order relationship has been debated [4–6]. As well, phylogenetic analysis using two ribosomal loci: the internal transcribed spacer region 1 (ITS1) and D1/D2 region of the large ribosomal subunit (D1/D2 LSU) has yielded multiple statistically-supported supra-genus groupings, although such topologies were often dependent on the locus examined, region amplified, taxa included in the analysis, and tree-building algorithm employed [7–9].

Therefore, while phenotypic and phylogenetic analyses suggest the existence of supra- genus relationships within the *Neocallimastigomycota*, the exact nature of such groupings is yet unclear. Approaches that utilize whole genomic and/or transcriptomic (henceforth referred to as –omics) datasets represent a promising tool towards resolving such relationships [10–14]. Comparative genomics approaches (e.g. calculation of shared Kmer (Kmer overlap) [15, 16], average nucleotide identity (ANI) [17], identification of genomic syntenic blocks [18]) have been increasingly utilized in taxonomic studies, aided by the development of lower cost high throughput sequencing technologies and the wider availability of bioinformatic analysis tools. More importantly, the development and implementation of phylogenomic approaches have been crucial in resolving high-rank [13], and intra-clade (e.g. [19]) phylogenies within fungi. Phylogenomic analysis involves the identification of groups of single-copy orthologous genes in the group of interest followed by individually multiple alignments of each orthologous gene.aligning such genes. Analysis to determine a species tree can then be performed on either the concatenated alignment of all genes to obtain a single phylogeny of the group in question, or on the individual alignments via coalescence of individual gene trees. In addition, the inferred gene trees canoutput from such approaches could also be compared to single gene phylogenies to assess their value and potential utility for taxonomic assessment and ecological surveys.

Within a Linnaean taxonomic framework, taxonomic associations between genera are accommodated in the intermediate ranks of families, orders, and classes. Currently, AGF genera are recognized in a single family (*Neocallimastigaceae*), order (*Neocallimastigales*), and class (*Neocallimastigomycetes*) in the phylum *Neocallimastigomycota* [20, 21]. It is interesting to note that a nomenclature novelty entry in Index Fungorum database (IF550425) proposes an additional family (*Anaeromycetacea*) with the genus *Anaeromyces* as its sole member, although no detailed justification for such a proposal was provided. Indeed, all current genera in the AGF, including *Anaeromyces*, are assigned to the family *Neocallimastigaceae* in recent publications [2, 3], reviews [4. 5.31-34 36], and databases (Mycobank, and Index Fungorum). Regardless, it is clear that the current intermediate rank taxonomic outline of AGF genera has not been proposed based on a detailed comparative phenotypic and phylogenetic analysis of relationships between genera. Rather, it reflects the cumulative and progressive recognition of the phylogenetic and phenotypic distinction of the Neocallimastigomycota when compared to all other fungal clades.

The earliest studies on AGF taxonomy [22] proposed accommodating them into a family (*Neocallimastigaceae*) within the chytrid order *Spizellomycetales*, a reflection of zoospore ultrastructure similarity; and emended the description of *Spizellomycetales* order to include zoopsores with multiple flagella. Ten years later, Li et al. [23] used cladistic analysis of 42 morphological and ultrastructural characters to demonstrate the distinction of the AGF when compared to members of the *Chytridiomycetes*, hence elevating the anaerobic gut fungi from a family to an order (Order *Neocallimastigales*). Molecular analysis using concatenated protein-coding genes as well as rRNA genes [21, 24, 25], and several morphological and ultrastructural differences from other *Chytridiomycetes* [26] necessitated their recognition as a phylum (*Neocallimastigomycota*) with one class (*Neocallimastigomycetes*), a view that has recently been corroborated via phylogenomic analysis [13]. Indeed, currently published taxonomic outlines, e.g. [20], and databases (e.g. GenBank [27], and Mycocosm [28]) recognize the AGF at the rank of phylum within the Mycota.

The last decade has witnessed a rapid expansion in the number of described genera within the *Neocallimastigomycota* [2, 4, 5, 29–34]. Due to such expansion, as well as the continuous recognition of the value of genome-based taxonomy in resolving relationships and circumscribing ranks in fungal taxonomy [10, 13, 14]; we posit that a lineage-wide phylogenomic assessment is warranted to resolve inter-genus relationships and explore the need for intermediate ranks to establish a proper Linnaean-based outline for the phylum. Here, we conducted transcriptomic sequencing on multiple additional AGF genera isolated and characterized in our laboratory, and combined these datasets with previously available AGF transcriptomes and genomes to resolve the inter-genus relationships within the *Neocallimastigomycota*. Based on our results, we propose accommodating AGF described genera into four distinct families to reflect the observed inter-genus relationships. In addition, we provide quantitative amino acid identity (AAI) for circumscribing such families, and test the utility of multiple single genes/loci as additional markers for resolving AGF phylogeny.

## Materials and Methods

### Cultures

Transcriptomes and genomes from fifty-two strains representing fourteen AGF genera were analyzed (Table 1). Of these, transcriptomes of twenty strains, representing six genera for which no prior sequence data were available were sequenced as part of this study. Many of the analyzed strains have previously been described as novel genera or species by the authors [5, 30- 32, 34] (Table 1). Others possessed identical features to previously described type strains and were designated as conferre (*cf.*) strains (Table 1). Few were identified to the genus level and given an alphanumeric strain name designation (Table 1).

**Table 1.** List of strains used in this study.

| Genus | species | Strain | Genome<br>BioProject<br>accession<br>number | Transcriptome<br>BioProject<br>accession<br>number | SRA accession<br>number | Assembled<br>transcriptome TSA<br>accession number | Reference |
| --- | --- | --- | --- | --- | --- | --- | --- |
| *Aestapascuomyces* | *dupliciliberans* | R1 |  | PRJNA847081 | SRR19612713 |  | This study |
| *Aklioshbomyces* | *papillarum* | WTS1 |  | PRJNA847081 | SRR19612712 |  | This study |
| *Anaeromyces* | *contortus* | ABS23 |  | PRJNA847081 | SRR19612701 |  | This study |
| *Anaeromyces* | *contortus* | C3G |  | PRJNA489922 |  | GGWR00000000.1 | [67, 68] |
| *Anaeromyces* | *contortus* | C3J |  | PRJNA489922 |  | GGWO00000000.1 | [67, 68] |
| *Anaeromyces* | *contortus* | G3G |  | PRJNA489922 |  | GGWP00000000.1 | [67, 68] |
| *Anaeromyces* | *contortus* | Na |  | PRJNA489922 |  | GGWN00000000.1 | [67, 68] |
| *Anaeromyces* | *contortus* | O2 |  | PRJNA489922 |  | GGWQ00000000.1 | [67, 68] |
| *Anaeromyces* | *mucronatus* | YE505 |  | PRJNA437872 |  |  | [38] |
| *Anaeromyces* | *robustus* | S4 | PRJNA330692 | PRJNA250973 |  |  | [69] |
| *Caecomyces* | *communis* | churrovis | PRJNA347164 | PRJNA393353 |  |  | [39, 41] |
| *Caecomyces* | *communis* | FD27 |  | PRJNA847081 | SRR19612700 |  | This study |
| *Caecomyces* | *communis* | TB33 |  | PRJNA847081 | SRR19612699 |  | This study |
| *Caecomyces* | *communis* | Iso3 |  | PRJNA489922 |  | GGXE00000000.1 | [67, 68] |
| *Caecomyces* | *communis* | Brit4 |  | PRJNA489922 |  | GGWS00000000.1 | [67, 68] |
| *Capellomyces* | *forminis* | Cap2a |  | PRJNA847081 | SRR19612698 |  | This study |
| *Cyllamyces* | *aberensis* | TSB2 |  | PRJNA847081 | SRR19612697 |  | This study |
| *Feramyces* | *austinii* | WSF2 |  | PRJNA489922 |  | GGWT00000000.1 | [67, 68] |
| *Feramyces* | *austinii* | WSF3 |  | PRJNA489922 |  | GGWU00000000.1 | [67, 68] |
| *Khyollomyces* | *ramosus* | ZO44 |  | PRJNA847081 | SRR19612696 |  | This study |
| *Liebetanzomyces* | *polymoprphus* | Orc37 |  | PRJNA847081 | SRR19612695 |  | This study |
| *Neocallimastix* | *frontalis* | EC30 |  | PRJNA847081 | SRR19612694 |  | This study |
| *Neocallimastix* | *frontalis* | Hef5 |  | PRJNA489922 |  | GGXJ00000000.1 | [67, 68] |
| *Neocallimastix* | *frontalis* | 27 |  | PRJNA437872 |  |  | [38] |
| *Neocallimastix* | *cameroonii* | G1 | PRJNA262392 | PRJNA251043 |  |  | [69] |
| *Neocallimastix* | *cameroonii* | lanati | PRJNA658393 | PRJNA677809 |  |  | [43] |
| *Neocallimastix* | *cameroonii* | G3 |  | PRJNA489922 |  | GGXC00000000.1 | [67, 68] |
| *Orpinomyces* | *joyonii* | AB6 |  | PRJNA847081 | SRR19612711 |  | This study |
| *Orpinomyces* | *joyonii* | AB3 |  | PRJNA847081 | SRR19612710 |  | This study |
| *Orpinomyces* | *joyonii* | ABC-24 |  | PRJNA847081 | SRR19612709 |  | This study |
| *Orpinomyces* | *joyonii* | D3A |  | PRJNA489922 |  | GGWV00000000.1 | [67, 68] |
| *Orpinomyces* | *joyonii* | D3B |  | PRJNA489922 |  | GGWW00000000.1 | [67, 68] |
| *Orpinomyces* | *joyonii* | D4C |  | PRJNA489922 |  | GGWX00000000.1 | [67, 68] |
| *Orpinomyces* | *joyonii* | SG4 |  | PRJNA437872 |  |  | [38] |
| *Paucimyces* | *polynucleatus* | BB3 |  | PRJNA847081 | SRR19612708 |  | This study |
| *Pecoramyces* | *ruminantium* | C1A | PRJNA200719 | PRJNA284193 |  |  | [67, 70] |
| *Pecoramyces* | *ruminantium* | S4B |  | PRJNA489922 |  | GGWY00000000.1 | [67, 68] |
| *Pecoramyces* | *ruminantium* | FS3C |  | PRJNA489922 |  | GGXF00000000.1 | [67, 68] |
| *Pecoramyces* | *ruminantium* | FX4B |  | PRJNA489922 |  | GGWZ00000000.1 | [67, 68] |
| *Pecoramyces* | *ruminantium* | YC3 |  | PRJNA489922 |  | GGXA00000000.1 | [67, 68] |
| *Pecoramyces* | *ruminantium* | Orc32 |  | PRJNA847081 | SRR19612707 |  | This study |
| *Pecoramyces* | *ruminantium* | AS31 |  | PRJNA847081 | SRR19612706 |  | This study |
| *Pecoramyces* | *ruminantium* | AS32 |  | PRJNA847081 | SRR19612705 |  | This study |
| *Pecoramyces* | *ruminantium* | F1 | PRJNA517297 | PRJNA517315 |  |  | [71] |
| *Piromyces* | *finnis* | finn | PRJNA330696 | PRJNA268530 |  |  | [69] |
| *Piromyces* | *finnis* | DonB11 |  | PRJNA847081 | SRR19612704 |  | This study |
| *Piromyces* | *cryptodigmaticus* | Axs23 |  | PRJNA847081 | SRR19612703 |  | This study |
| *Piromyces* | *cryptodigmaticus* | A1 |  | PRJNA489922 |  | GGXB00000000.1 | [67, 68] |
| *Piromyces* | *potentiae* | B4 |  | PRJNA489922 |  | GGXH00000000.1 | [67, 68] |
| *Piromyces* | *sp. NZB19* | Ors32 |  | PRJNA847081 | SRR19612702 |  | This study |
| *Piromyces* | *sp. PR1* | E2 | PRJNA82799 |  |  |  | [69] |
| *Piromyces* | *rhizinflatus* | YM600 |  | PRJNA437872 |  |  | [38] |

### RNA extraction, Sequencing, quality control, and transcripts assembly

Isolates were grown in rumen fluid medium with cellobiose as the sole carbon source [35] to late log/early stationary-phase (approximately 48 to 60 h post inoculation). Cultures were vacuum filtered to obtain fungal biomass then grounded with a pestle under liquid nitrogen. Total RNA was extracted using Epicentre MasterPure yeast RNA purification kit (Epicentre, Madison, WI) according to manufacturer’s instructions and stored in RNase-free Tris-EDTA buffer. Transcriptomic sequencing using Illumina HiSeq2500 platform and 2L×L150 bp paired-end library was conducted using the services of a commercial provider (Novogene Corporation, Beijing, China), or at the Oklahoma State University Genomics and Proteomics center. The RNA-seq data were quality trimmed and *de novo* assembled with Trinity (v2.6.6) using default parameters. For each data set, redundant transcripts were clustered using CD-HIT [36] with identity parameter of 95% (–c 0.95). The obtained nonredundant transcripts were subsequently used for peptide and coding sequence prediction using the TransDecoder (v5.0.2) (https://github.com/TransDecoder/TransDecoder) with a minimum peptide length of 100 amino acids. Assessment of transcriptome completeness per strain was conducted using BUSCO [37] with the fungi_odb10 dataset modified to remove 155 mitochondrial protein families as previously suggested [38].

### Phylogenomic analysis

The phylogenomic analysis includes 20 newly sequenced and 32 existing AGF genomic and transcriptome sequences (Table 1) [38–43]. Five *Chytridiomycota* genomes were also included as the outgroup (*Chytriomyces* sp. strain MP 71, *Entophlyctis helioformis* JEL805, *Gaertneriomyces semiglobifer* Barr 43, *Gonapodya prolifera* JEL478, and *Rhizoclosmatium globosum* JEL800 [44, 45]). The “fungi_odb10” dataset including 758 phylogenomic markers for Kingdom Fungi was retrieved from BUSCO v4.0 package, and used in our analysis. Profile hidden-Markov-models of these markers were created and used to identify homologues in all included fifty-eight fungal proteomes using hmmer3 (v3.1b2) employed in the PHYling pipeline (https://doi.org/10.5281/zenodo.1257002). A total of 670 out of the 758 “fungi_odb10” markers were identified with conserved homologs in the 57 AGF and Chytrids genomes, which were then aligned and concatenated for the subsequent phylogenomic analyses. The final input data include 491,301 sites with 421,690 distinct patterns. The IQ-TREE v1.7 package was used to find the best-fit substitution model and reconstruct the phylogenetic tree with the maximum-likelihood approach.

### Average amino acid identity

We calculated Average Amino acid Identity (AAI) values for all possible pairs in the dataset using the predicted peptides output from TransDecoder.LongOrfs. AAI values were generated using the aai.rb script available as part of the enveomics collection [46]. Through reciprocal all versus all protein Blast, AAI values represent indices of pairwise genomic relatedness [47]. Since its introduction in 2005 [47] as a means for standardizing taxonomy at ranks higher than species, AAI has been extensively used in bacterial and archaeal genome-based taxonomic studies [48–50]. However, AAI has been utilized only sparsely in the fungal world (e.g. [51, 52], with genome-based quantitative comparisons (e.g. Jaccard index of genomic distance (the fraction of shared k-mers), identification of syntenic blocks, and Average Nucleotide Identity (ANI) [15, 18]) being more heavily utilized and often for delineating lower taxonomic level (e.g. species) boundaries. AAI, however, has the advantage of being readily conducted on the predicted peptides from transcriptomic datasets, as it uses amino acid sequences. The ease of obtaining transcriptomic rather than genomic sequences for AGF (mostly due to the extremely high AT content in intergenic regions and the extensive proliferation of microsatellite repeats, often necessitating employing multiple sequencing technologies for successful genomic assembly) makes the use of AAI for delineation of taxonomic boundaries more appealing.

### Single gene phylogenetic analysis

Two ribosomal loci (D1/D2 LSU, and ITS1) and four protein-coding gene trees (RNA polymerase II large subunit (RPB1), RNA polymerase II second largest subunit (RPB2), Minichromosome maintenance complex component 7 (MCM7), and Elongation factor 1-alpha (EF1α)) were evaluated. Sequences for ITS1 and D1/D2 LSU were either obtained from prior studies [5, 9, 30–32, 34, 53] or were bioinformatically extracted from genomic assemblies [54]. Amino acids sequences of RPB1, RPB2, MCM7 and EF1α were obtained from the *Anaeromyces robustus* genome (GenBank assembly accession number: GCA_002104895.1), and used as bait for Blastp searches against all predicted proteomes in all transcriptomic datasets. Sequences for each protein, as well as for the rRNA loci were aligned using MAFFT with default parameters. The alignments were used as inputs to IQ-TREEtree [55, 56] first to predict the best substitution model (using the lowest BIC criteria) and to generate maximum likelihood trees under the predicted best model. The “–alrt 1000” option for performing the Shimodaira-Hasegawa approximate-likelihood ratio test (SH-aLRT), “-abayes” option for performing approximate Bayes test, and the “–bb 1000” option for ultrafast bootstrap (UFB) were added to the IQ-TREE command line, which resulted in the generation of phylogenetic trees with three support values (SH-aLRT, aBayes, and UFB) on each branch.

### Nucleotide sequencing accession number

Raw Illumina RNA-seq read sequences are deposited in GenBank under the BioProject accession number PRJNA847081 and BioSample accessions numbers SAMN28920465- SAMN28920484. Individual SRA accessions are provided in Table 1.

## Results

### Sequencing

Transcriptomic sequencing yielded 15.6 to 23.8 (average 19.82) million reads that were assembled into 22,649 to 106,687 total transcripts, 20,599 to 103,405 distinct transcripts (clustering at 95%; average 40,099), and 13,858 to 28,405 predicted peptides (average 19,667) (Table S2). Assessment of transcriptome completion using BUSCO yielded high values (73.63 to 99.5%) for all assemblies (Table S1).

**Table 2.** Clades circumscribed in this study.

| Clades | Genera | AAI |  |  | Phenotype |
| --- | --- | --- | --- | --- | --- |
|  |  | Average intra-genus (range) | Average inter-genus intra-clade (range) | Average inter-clade (range) |  |
| Clade 1 | *Pecoromyces*, *Orpinomyces*, *Neocallimastix*, *Aestipascuomyces*, *Feromyces* | 96.89 (87.78-99.49) | 82.95 (75.44-78.99) | 73.21 (65.27-76.64) | Polyflagellated zoospores except for *Pecoromyces* |
| Clade 2 | *Cyllamyces*, *Caecomyces* | 94.01 (88.02-98.37) | 84.05 (83.08-83.67) | 72.8 (67.39-74.91) | Bulbous rhizoidal growth pattern |
| Clade 3 | *Piromyces* | 79.35 (72.58-99.06) | 79.35 (72.58-99.06) | 72.61 (65.27-75.61) | Monocentric thalli, monoflagellated zoospores, filamentous rhizoidal growth pattern |
| Clade 4 | *Anaeromyces*, *Liebetanzomyces*, *Capellomyces* | 96.55 (93.07-99.6) | 84.41 (83.58-85.48) | 73.75 (67.25-76.64) | Filamentous rhizoidal growth pattern, monoflagellated zoospore, all monocentric thallus except *Anaeromyces* |
|  | *Aklioshbomyces* | NA | NA | 73.54 (69.1-75.42) |  |
|  | *Paucimyces* | NA | NA | 74.26 (68.14-76.98) |  |
|  | *Khyollomyces* | NA | NA | 71.88 (66.47-73.41) |  |

### Resolving inter-genus relationships in the Neocallimastigomycota

Multiple supra-genus relationships were well supported in all phylogenomic outputs. Four distinct clades were observed (Figure 1 and Table 2). Clade one constituted members of the genera *Pecoramyces, Orpinomyces, Neocallimastix, Feramyces* and *Aestipascuomyces.* Within this large clade, a strong support for *Pecoramyces* and *Orpinomyces* association, as well as for *Neocallimastix*, *Aestipascuomyces, and Feramyces* association was observed (Figure 1). Phenotypically, this clade encompasses all the AGF genera producing polyflagellated zoospores; and all members of the clade, with the exception the genus *Pecoramyces* produce polyflagellated zoospores. Clade two constituted members of the genera *Cyllamyces* and *Caecomyces.* Phenotypically, this clade encompasses the two genera exhibiting a bulbous rhizoidal growth pattern in the *Neocallimastigomycota*. Clade three constituted members of the genus *Piromyces*. Compared to all other AGF genera, the genus *Piromyces* currently exhibits high intra-genus sequence divergence based on ITS1 and LSU analysis [3]. The genus was first defined to encompass all phenotypes with monocentric thalli, filamentous rhizoidal system, and monoflagellated zoospores [57]. However, subsequent isolation efforts clearly demonstrated that such phenotype is prevalent in a wide range of phylogenetically disparate genera across the *Neocallimastigomycota* [4, 5, 29]. Currently, *Piromyces* encompasses all taxa phylogenetically affiliated with the first described monocentric, monoflagellated, and filamentous isolate (*Piromyces communis* [57]). Clade four constituted members of the genera *Anaeromyces*, *Liebetanzomyces*, and *Capellomyces*. The clade encompasses genera with filamentous rhizoidal system, and monoflagellated zoospores. The genus *Anaeromyces* produces polycentric thalli, while the genera *Liebetanzomyces*, and *Capellomyces* produce monocentric thalli.

**Figure 1.**
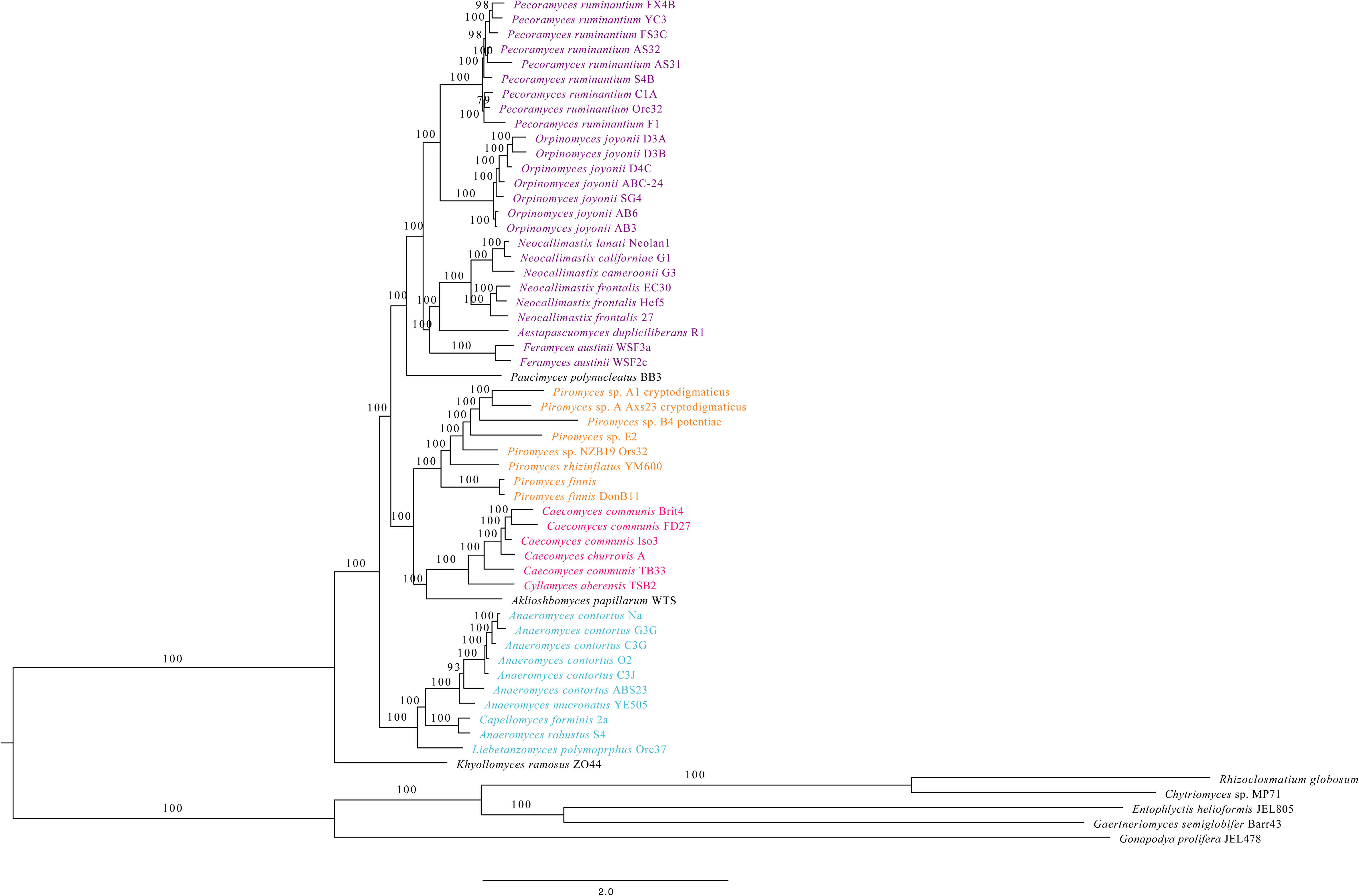
Phylogenomic tree of *Neocallimastigomycota* based on 670 genome-wide markers highlighting the family-level relationships within the phylum. The tree was reconstructed using the maximum likelihood approach implemented in the IQ-TREE package. Number on each branch represents the ultrafast bootstrap value suggesting the robustness of the taxa joining. The scale bar at the bottom indicates the number of substitutions per site in the analysis. Isolate names at tree tips are color coded by clade (clade 1, purple; clade 2, lavender; clade 3, orange; clade 4, light blue).

Few genera clustered outside these four clades described above. The genera *Paucimyces* and *Aklioshbomyces* formed distinct branches at the base of clades 1 and 2, respectively (Figure 1). Finally, the position of the genus *Khoyollomyces* was unique and solitary, being consistently located at the base of the tree, suggesting its deep-branching and relatively ancient origin.

### Estimating AAI identities

AAI values were estimated using the entire dataset of predicted peptides (Figure 2). Intra-genus AAI values ranged between 72.58-99.6% (Average 92.16 ± 8.55). However, the low intra-genus divergence estimates were only confined to the broadly circumscribed genus *Piromyces*. Indeed, excluding *Piromyces* from this analysis, intra-genus AAI values ranged between 87.78-99.6%, (Average 95.67 ± 3.41). Pairwise AAI values for members of different genera within the same clade (intra-clade inter-genus AAI values) ranged between 75.44-85.48% (Average 79.58 ± 2.47). Maximum intra-clade inter-genus divergence was observed between members of the genera *Neocallimastix* and *Pecoramyces* (Average 77.5 ± 0.91) and the genera *Neocallimastix* and *Orpinomyces* (Average 77.4 ± 0.59) in clade 1, while minimal intra-clade inter-genus divergence were observed between *Caecomyces* and *Cyllamyces* in clade 2 (83.7% ± 0.4); as well as the genera *Anaeromyces* and *Capellomyces* (Average 84.5 ± 0.57), the genera *Anaeromyces* and *Liebetanzomyces* (Average 83.9 ± 0.3), and the genera *Capellomyces* and *Liebetanzomyces* (Average 85.1 ± 0.18) in clade 4. Inter-clade AAI values averaged 73.15 ± 1.57, and ranged between 65.27% (between members of the genera *Piromyces* and *Neocallimastix*) and 76.64 % (between members of the genera *Capellomyces* and *Pecoramyces*).

**Figure 2.**
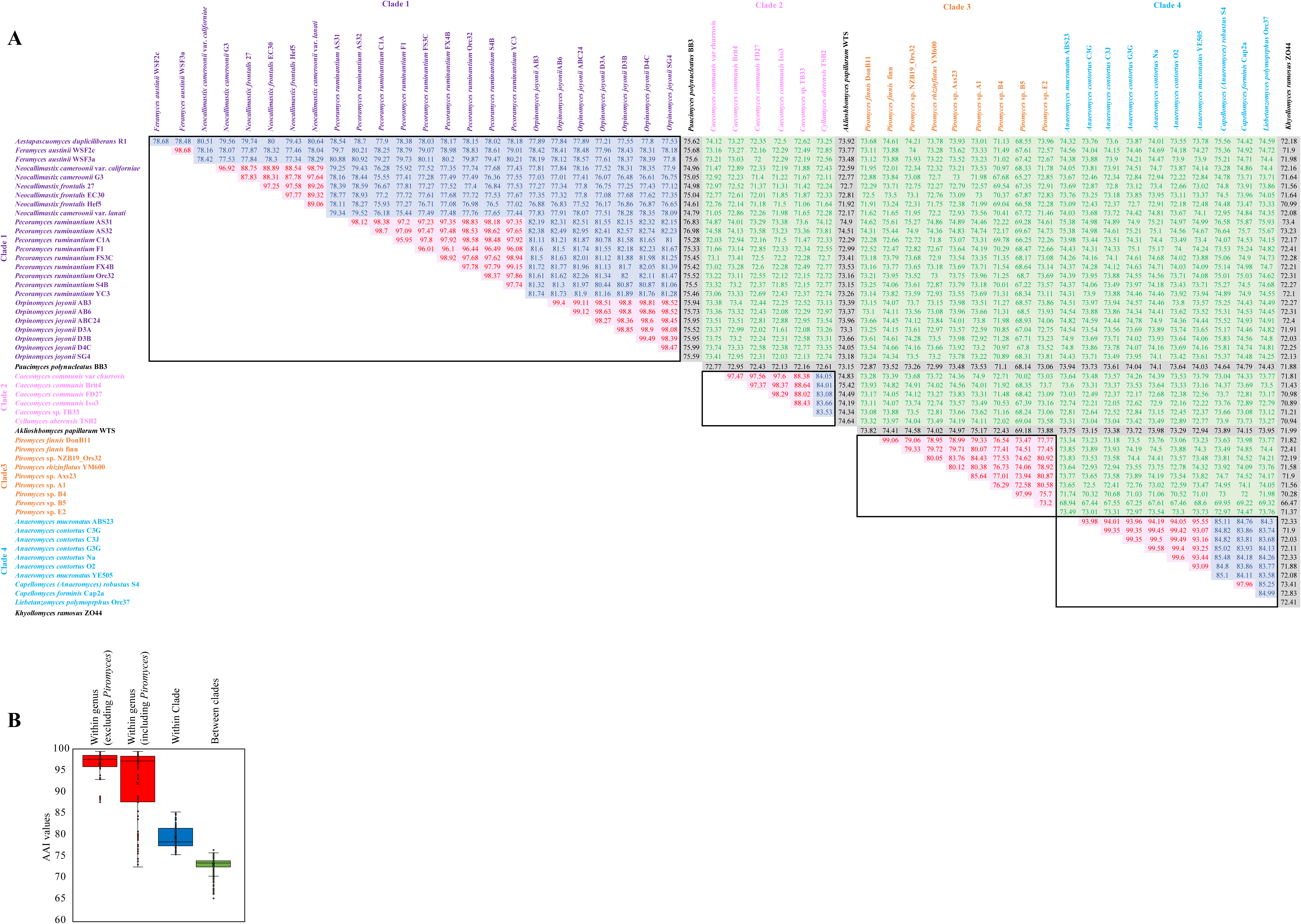
Upper triangle matrix (A) and box and whisker plots (B) for the AAI values obtained for all possible pairwise comparisons of the datasets analyzed in this study. (A) Isolate names in rows and columns are color coded by clade (clade 1, purple; clade 2, lavender; clade 3, orange; clade 4, light blue). The AAI values for each clade are shown within a thick border. Intra-genus values are shown in red text with pink highlight, intra-clade/ inter-genus values are shown in blue text with light blue highlight, while inter-clade values are shown in green text with light green highlight. Values for the three genera unaffiliated with the 4 clades are highlighted in grey. (B) Box and whisker plots constructed using the values in (A). Intra-genus values (red) are shown both including and excluding the genus *Piromyces*. Intra-clade/ inter-genus values are shown in blue. Inter-clade values are shown in green. Each box plot spans the region between the 25-percentile to 75-percentile, while the whiskers limit the minimum and maximum scores excluding the outliers. The thick line inside the box marks the median, while the ‘x’ corresponds to the average value.

### Single gene phylogenetic analysis for resolving AGF inter-genus relationships

We tested whether supra-genus clades topology as well as within clades inter-genus relationships observed in phylogenomic analysis were retained in single gene phylogenies (Figure 3-8). One ribosomal locus (D1/D2 LSU) and one protein-coding gene (RPB1) retained the monophylly of all four clades described above (Figure 3, 5, Table S2). As well, both D1/D2 LSU and RPB1 phylogenies resolved all inter-genus relationships within all clades in the *Neocallimastigomycota* (Figure 3, 5). On the other hand, ITS1, RPB2, MCM7, and EF1α phylogenies each recovered three out of the four supra-genus clades delineated above. The monophylly of clade 1 was not retained in ITS1 and RPB2 phylogenies (Figure 4, 6, Table S2), the monophylly of clade 4 was not retained in MCM7 phylogeny (Figure 7), and the monophylly of clade 3 was not retained in EF1α phylogeny (Figure 8). Further, within the clades that were supported, few inter-genus relationships were compromised in ITS1 (genus *Anaeromyces*), and EF1α (genera *Neocallimastix, Orpinomyces,* and *Pecoramyces*) phylogenies.

**Figure 3.**
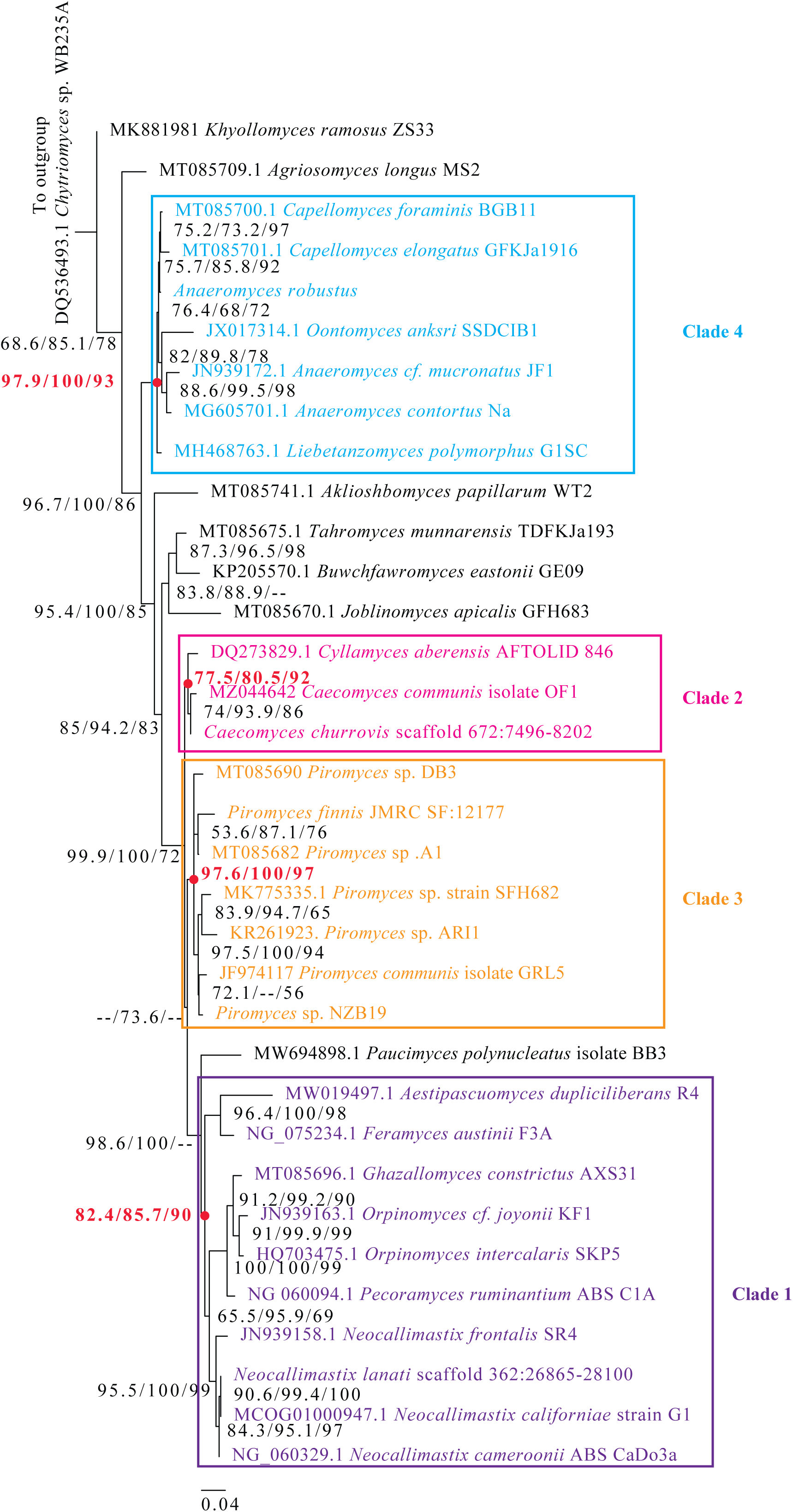
Maximum likelihood phylogenetic tree constructed using the D1/D2 region of the LSU rRNA genes of all cultured and described *Neocallimastigomycota* genera. Sequences were either obtained from prior studies [5, 9, 30–32, 34, 53] or were bioinformatically extracted from genomic assemblies [54], and GenBank accession numbers are shown for each branch label. Sequences were aligned using MAFFT with default parameters. IQ-tree [55, 56] was used to choose the best substitution model (TN+F+G4 was chosen using the lowest BIC criteria) and to generate the maximum likelihood tree. Support values at each node correspond to SH-aLRT, aBayes, and ultrafast bootstrap. Clades are coded using the same color code in Figure 2 (clade 1, purple; clade 2, lavender; clade 3, orange; clade 4, light blue), and boxes with the same colors are used to delimit each clade. The support values at the nodes corresponding to each clade are shown in bold red text, and the node itself is shown as a red dot. The tree was rooted (root not shown) using the D1/D2 region of the LSU rRNA gene from *Chytriomyces* sp. WB235A (GenBank accession number DQ536493.1).

**Figure 4.**
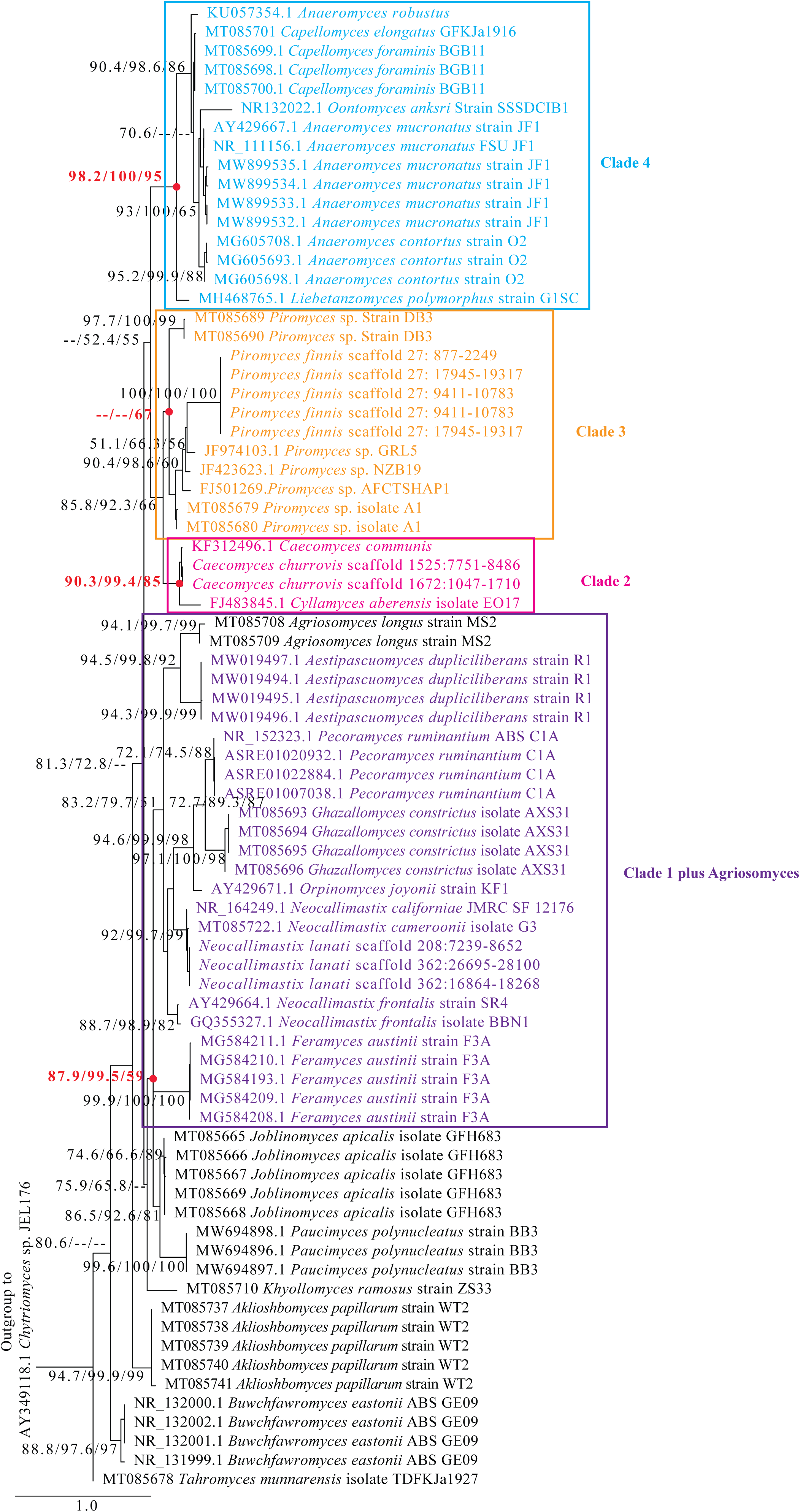
Maximum likelihood phylogenetic tree constructed using the ITS1 region of all cultured and described Neocallimastigomycota genera. Sequences were either obtained from prior studies [5, 9, 30–32, 34, 53] or were bioinformatically extracted from genomic assemblies [54], and GenBank accession numbers are shown for each branch label. Sequences were aligned using MAFFT with default parameters. IQ-tree [55, 56] was used to choose the best substitution model (TN+F+G4 was chosen using the lowest BIC criteria) and to generate the maximum likelihood tree. Support values at each node correspond to SH-aLRT, aBayes, and ultrafast bootstrap. Branch labels are color coded using the same color code in Figure 2 (clade 1, purple; clade 2, lavender; clade 3, orange; clade 4, light blue), and boxes with the same colors are used to delimit each clade. The support values at the nodes corresponding to each clade are shown in bold red text, and the node itself is shown as a red dot. The tree was rooted (root not shown) using the ITS1 region from *Chytriomyces* sp. JEL176 (GenBank accession number AY349118.1).

**Figure 5.**
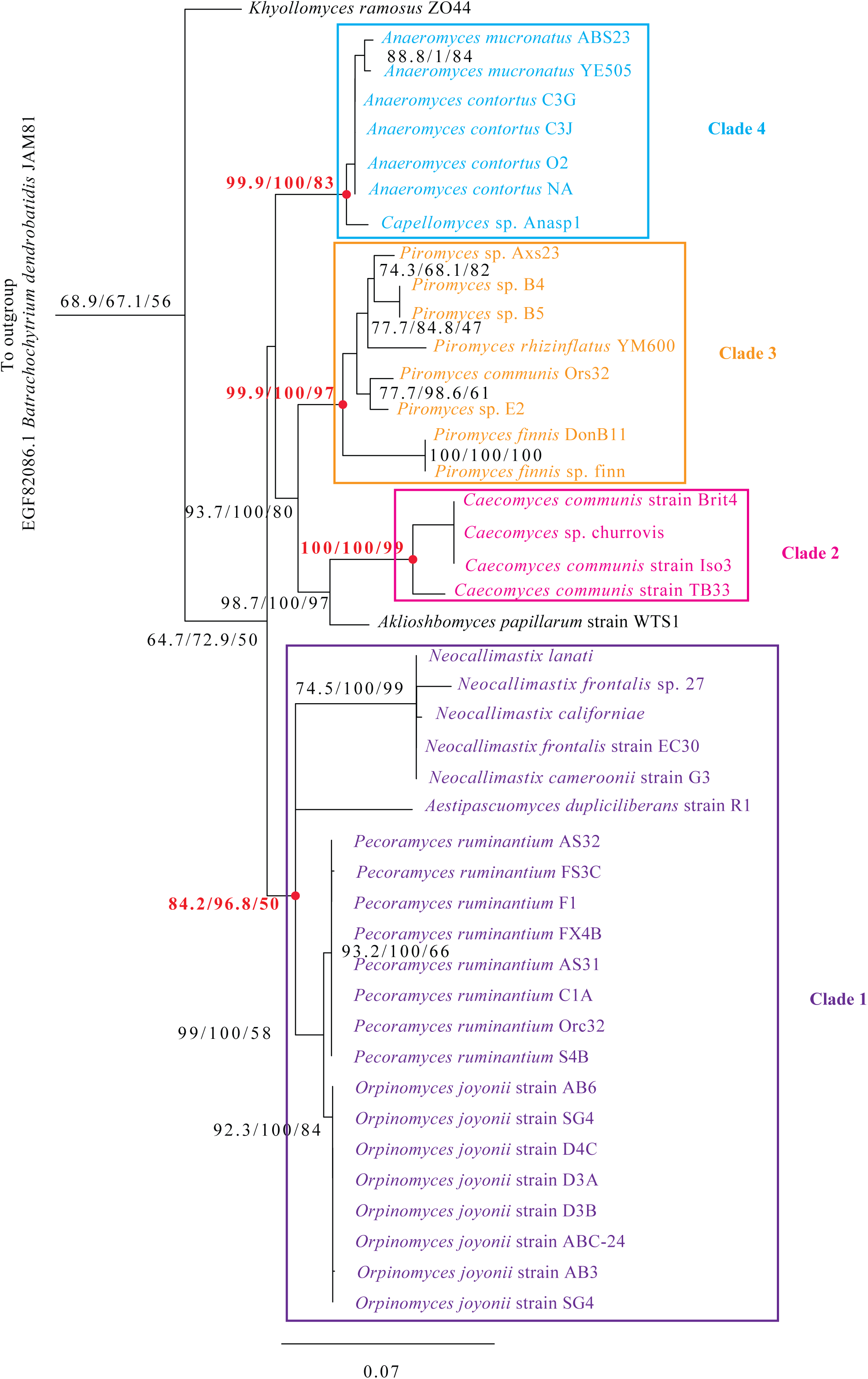
Maximum likelihood phylogenetic tree constructed using the protein sequences of the largest subunit of DNA-dependent RNA polymerase II (RPB1). Amino acids sequence of RPB1 was obtained from the *Anaeromyces robustus* genome (GenBank assembly accession number: GCA_002104895.1), and used as bait for Blastp searches against all predicted proteomes in all transcriptomic datasets. Sequences were aligned using MAFFT with default parameters. IQ-tree [55, 56] was used to choose the best substitution model (LG+R2 was chosen using the lowest BIC criteria) and to generate the maximum likelihood tree. Support values at each node correspond to SH-aLRT, aBayes, and ultrafast bootstrap. Branch labels are color coded using the same color code in Figure 2 (clade 1, purple; clade 2, lavender; clade 3, orange; clade 4, light blue), and boxes with the same colors are used to delimit each clade. The support values at the nodes corresponding to each clade are shown in bold red text, and the node itself is shown as a red dot. The tree was rooted (root not shown) using the RPB1 sequence from *Batrachochytrium dendrobatidis* JAM81 (GenBank accession number EGF82086.1).

**Figure 6.**
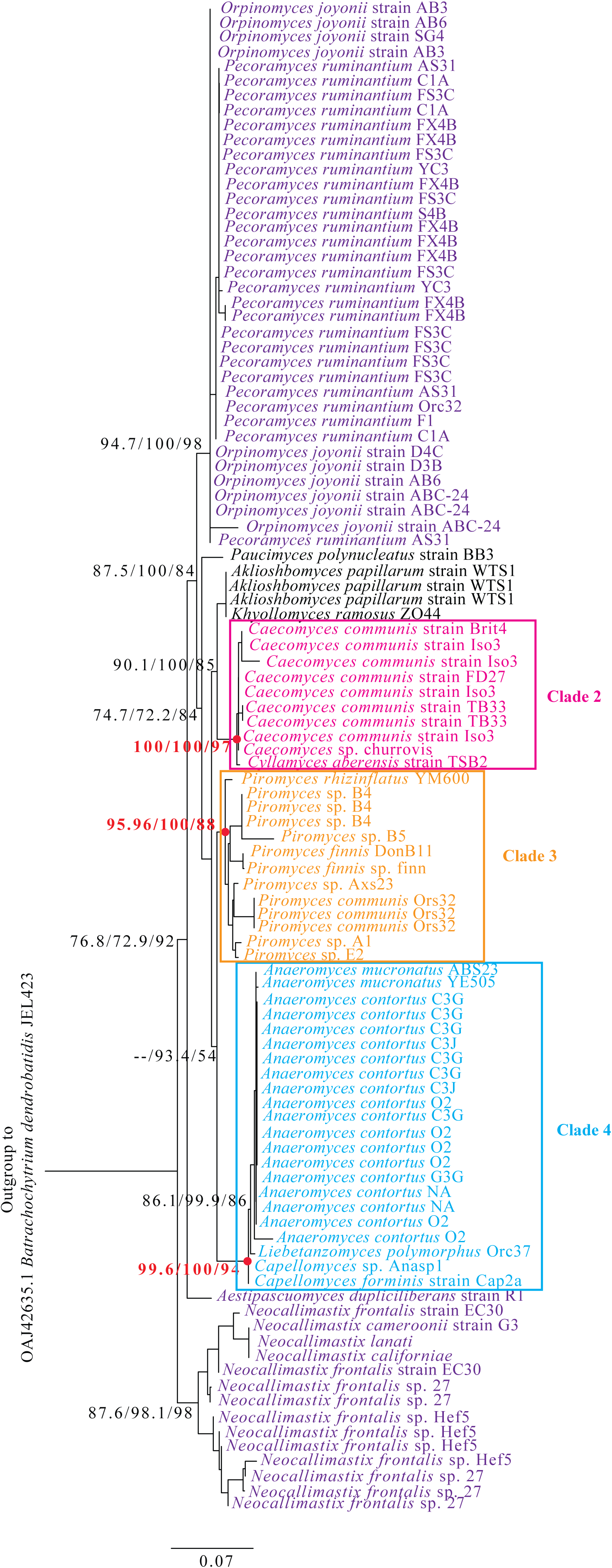
Maximum likelihood phylogenetic tree constructed using the protein sequences of the second largest subunit of DNA-dependent RNA polymerase II (RPB2). Amino acids sequence of RPB2 was obtained from the *Anaeromyces robustus* genome (GenBank assembly accession number: GCA_002104895.1), and used as bait for Blastp searches against all predicted proteomes in all transcriptomic datasets. Sequences were aligned using MAFFT with default parameters. IQ-tree [55, 56] was used to choose the best substitution model (LG+R3 was chosen using the lowest BIC criteria) and to generate the maximum likelihood tree. Support values at each node correspond to SH-aLRT, aBayes, and ultrafast bootstrap. Branch labels are color coded using the same color code in Figure 2 (clade 1, purple; clade 2, lavender; clade 3, orange; clade 4, light blue), and boxes with the same colors are used to delimit each clade if the clade is supported. The support values at the nodes corresponding to each clade are shown in bold red text, and the node itself is shown as a red dot. The tree was rooted (root not shown) using the RPB2 sequence from *Batrachochytrium dendrobatidis* JEL423 (GenBank accession number OAJ42635.1).

**Figure 7.**
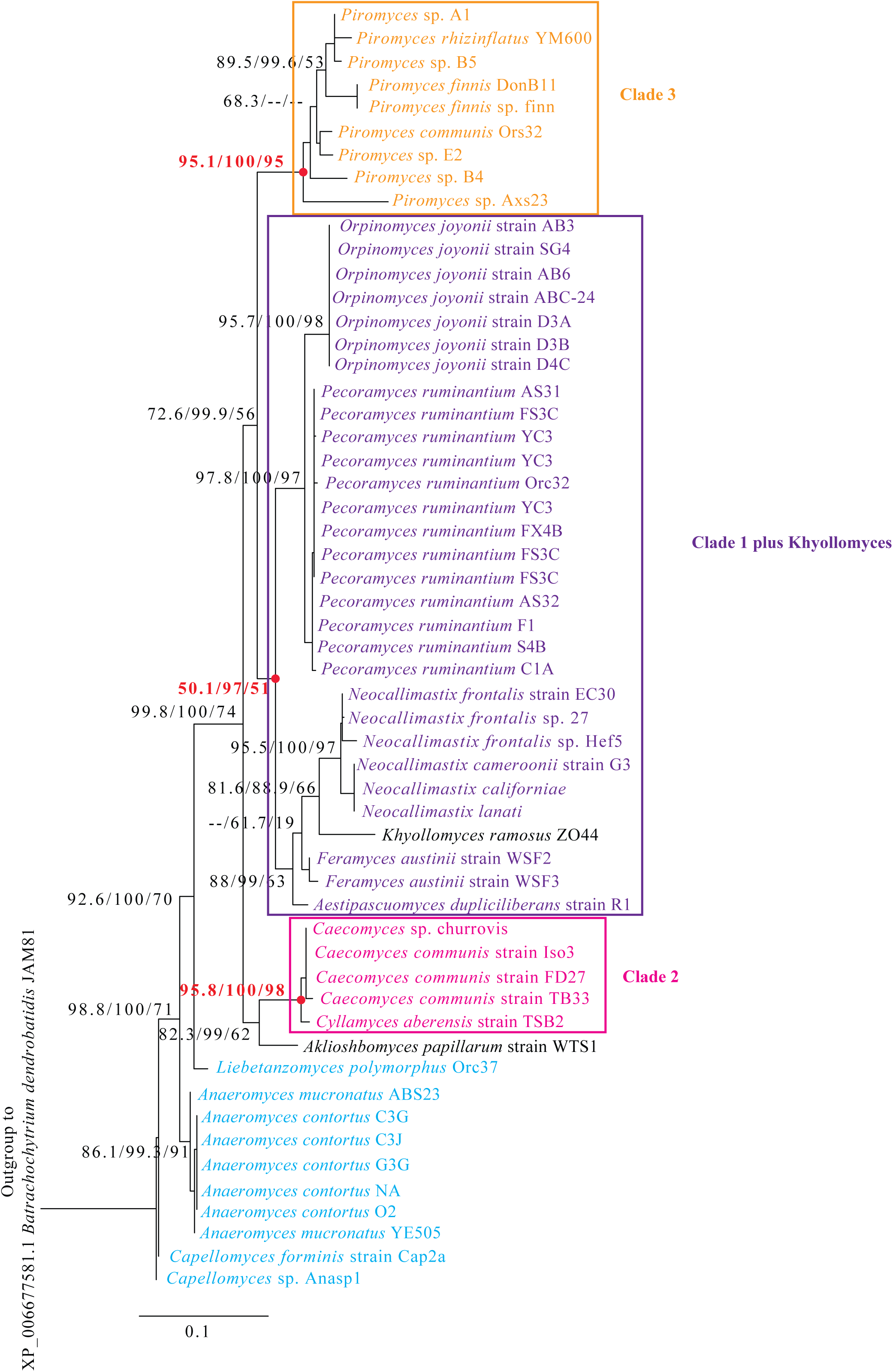
Maximum likelihood phylogenetic tree constructed using the protein sequences of the DNA replication licensing factor MCM7. Amino acids sequence of MCM7 was obtained from the *Anaeromyces robustus* genome (GenBank assembly accession number: GCA_002104895.1), and used as bait for Blastp searches against all predicted proteomes in all transcriptomic datasets. Sequences were aligned using MAFFT with default parameters. IQ-tree [55, 56] was used to choose the best substitution model (LG+R3 was chosen using the lowest BIC criteria) and to generate the maximum likelihood tree. Support values at each node correspond to SH-aLRT, aBayes, and ultrafast bootstrap. Branch labels are color coded using the same color code in Figure 2 (clade 1, purple; clade 2, lavender; clade 3, orange; clade 4, light blue), and boxes with the same colors are used to delimit each clade if the clade is supported. The support values at the nodes corresponding to each clade are shown in bold red text, and the node itself is shown as a red dot. The tree was rooted (root not shown) using the MCM7 sequence from *Batrachochytrium dendrobatidis* JAM81 (GenBank accession number XP_006677581.1).

**Figure 8.**
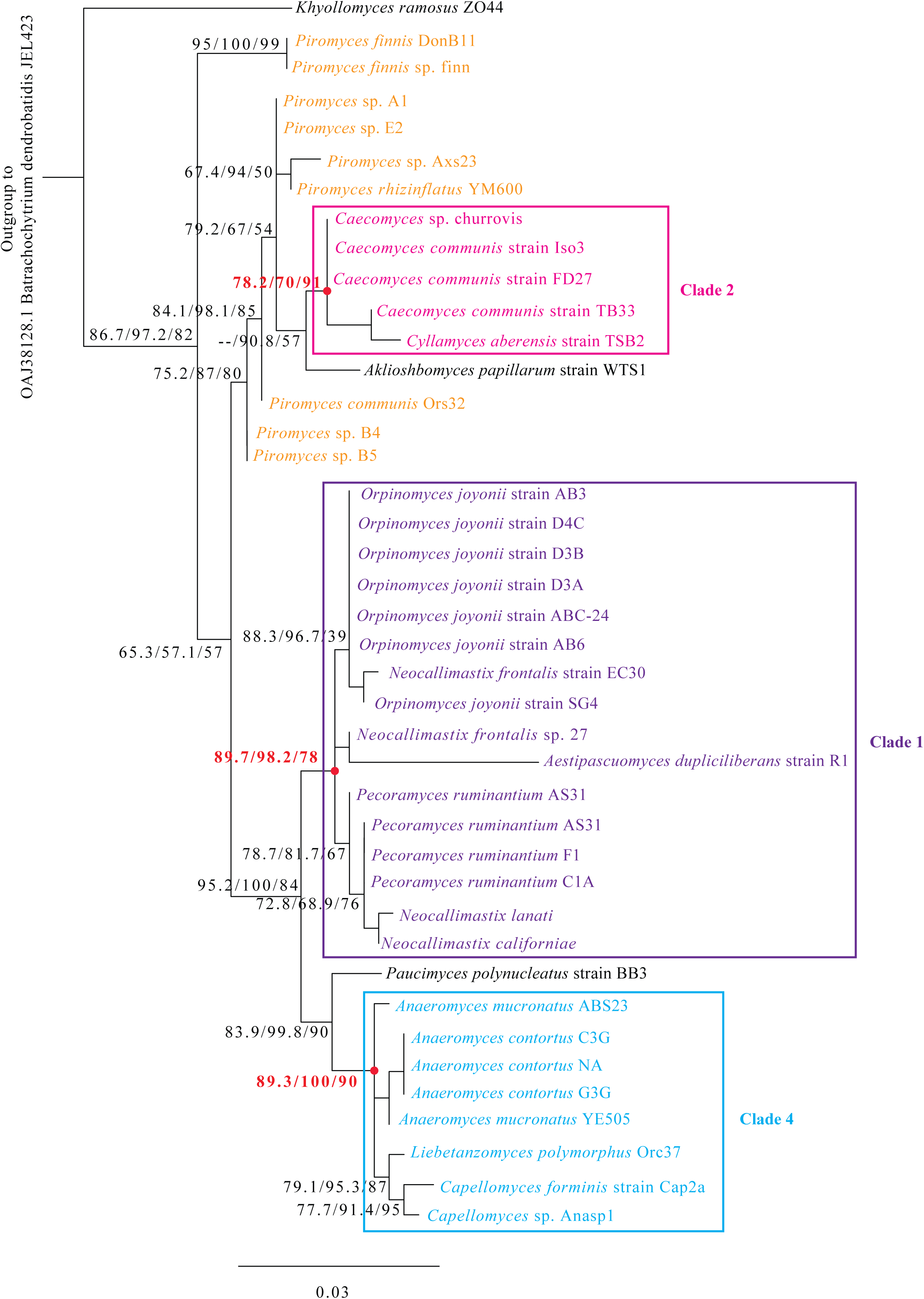
Maximum likelihood phylogenetic tree constructed using the protein sequences of the elongation factor 1-alpha (EF-1A). Amino acids sequence of EF-1A was obtained from the *Anaeromyces robustus* genome (GenBank assembly accession number: GCA_002104895.1), and used as bait for Blastp searches against all predicted proteomes in all transcriptomic datasets. Sequences were aligned using MAFFT with default parameters. IQ-tree [55, 56] was used to choose the best substitution model (LG+R2 was chosen using the lowest BIC criteria) and to generate the maximum likelihood tree. Support values at each node correspond to SH-aLRT, aBayes, and ultrafast bootstrap. Branch labels are color coded using the same color code in Figure 2 (clade 1, purple; clade 2, lavender; clade 3, orange; clade 4, light blue), and boxes with the same colors are used to delimit each clade if the clade is supported. The support values at the nodes corresponding to each clade are shown in bold red text, and the node itself is shown as a red dot. The tree was rooted (root not shown) using the EF-1A sequence from *Batrachochytrium dendrobatidis* JEL423 (GenBank accession number OAJ38128.1).

## Discussion

### Identifying and circumscribing supra-genus relationships within the Neocallimastigomycota

Our phylogenomic analysis identified four distinct statistically supported supra-genus clades in the *Neocallimastigomycota* (Table 2, Figure 1). Clades’ boundaries were based on phylogenomic tree topologies, while taking taxonomically informative morphological characteristics into account. For example, phylogenomic analyses placed the genus *Paucimyces* at the base of clade 1, and the genus *Aklioshbomyces* at the base of clade 2. Exclusion of *Paucimyces* from clade 1 was based on its production of monoflagellated zoospores [32], as opposed to the polyflagellated zoospores produced by all members of clade 1 (with the exception of *Pecoramyces*). Similarly, exclusion of *Aklioshbomyces* from clade 2 was based on its filamentous rhizoidal growth pattern; which contrasts the bulbous growth pattern exclusive to both genera (*Caecomyces* and *Cyllamyces*) constituting clade 2.

AAI values were further examined to quantitatively circumscribe these clades. A clear delineation of the clade boundary was evident using AAI values (Figure 2). Within genus, AAI values ranged between 87.78-99.6% (or 72.58-99.6% if including values for the broadly circumscribed genus *Piromyces*). Inter-genus/ Intra-clade AAI estimates ranged between 75.44- 85.48%, while inter-clade values ranged between 65.27-76.64% (Figure 2). These values are similar to AAI values estimated for delineating the *Ascomycetes* family *Hypoxylaceae* [51], but are higher than the arbitrary cutoffs used for delineating taxa in the prokaryotic world (∼45-65% for family, ∼65-95% for genus [48]). Therefore, we suggest using 85.0%, and 75.0% AAI cutoff values as a guide for circumscribing genera, and families, respectively, in the *Neocallimstigomycota*. Currently, the genus *Piromyces* represents the sole genus in clade 3. AAI estimates using the currently available *Piromyces* species –omics datasets suggest broader inter-genus AAI range when compared to other genera (Figure 2). This is a reflection of the fact that the genus was originally circumscribed based on phenotypic, rather than a combination of phenotypic and molecular data. Future availability of additional –omics data coupled to a detailed comparative morphotypic analysis of its described species could possibly lead to splitting this genus (the sole member of clade 3 here) into several clades.

Up to this point, only ITS1 and D1/D2 LSU loci have been evaluated for assessment of phylogenetic positions of genera within the Neocallimastigomycota, as well as for ecological culture-independent surveys [7, 9]. To test the utility of other phylomarkers commonly utilized in fungal taxonomy, we assessed additional four protein-coding genes, and examined concordance between each of the six loci (ribosomal ITS1 and D1/D2 LSU, and RPB1, RPB2, MCM7, and EF-1α) and phylogenomic trees topologies. Our results demonstrate that D1/D2 LSU, currently regarded as the phylomarker of choice for genus-level delineation [9, 58] and utilized as a marker in culture-independent diversity surveys [9], is equally useful in resolving supra-genus clades delineated by phylogenomics (Figures 3, S1). As well, our results add the protein-coding gene RPB1 to the list of phylomarkers that could be used for inter-genus, and supra-clade delineation (Figures 5, S2). As such, values of 8.5%, and 2.1% for LSU, and RPB1, respectively (these values correspond to the 75-percentile value for intra-clade inter-genus divergence based on the distance matrix from the alignments used to generate the maximum likelihood trees in Figures 3, 5) seem to circumscribe these clades. The high sequence similarity in the protein-coding gene RPB1 is quite surprising since, typically, higher levels of divergence are usually observed in protein coding genes when compared to the non-protein-coding ribosomal genes [59]. Other phylomarkers tested here were only successful in resolving three of the four clades, and some also compromised intra- and inter-genus relationships (Figures 4, 6-7).

Such failure to resolve genus-level relationships appears to be a function of high sequence similarities in these genes. For example, the inter-genus divergence values between *Orpinomyces* and *Pecoramyces* RPB2 sequences ranged between 0-1.8%, which are comparable to the values within the genus *Orpinomyces*. This has resulted in failure of RPB2 to resolve the *Orpinomyces*- *Pecoramyces* relationship. The unreliability of the ITS1 locus for clade delineation has been described before, and is mainly due to length variability between genera and high within-strain sequence divergence [7, 9].

### Phylogenetic position of taxa currently lacking genome or transcriptome sequences

The fifty-three transcriptomic datasets examined cover fourteen out of the twenty currently described AGF genera. The remaining six genera (*Oontomyces*, *Buwchfawromyces, Agriosomyces, Ghazallomyces, Tahromyces,* and *Joblinomyces*) are all currently represented by a single species. Further, most of these genera appear to exhibit extremely limited geographic and animal host distribution patterns [4, 5, 9, 29]. The phylogenetic position of these six genera could hence be only evaluated using available D1/D2 LSU (and to some extent ITS1) sequence data from taxa description publications. D1/D2 LSU and ITS1 phylogenies strongly support placement of the genus *Ghazallomyces* as a member of clade 1 (Figure 3, 4) [5]. Further, the genus produces polyflagellated zoospores (an exclusive trait for clade 1), filamentous rhizoid (similar to all taxa in clade 1), and monocentric thalli (similar to all genera in clade 1, except *Orpinomyces*), further supporting its recognition as member of clade 1[5]. Similarly, phylogenetic analysis using D1/D2 LSU and ITS1 supports the placement of genus *Oontomyces* as a member of clade 4 (Figure 3, 4). Members of the genus *Oontomyces* exhibit similar phenotypes (monocentric thalli, monoflagellated zoospores, and filamentous rhizoidal growth patterns) to the genera *Liebetanzomyces* and *Capellomyces* in the clade [29].

Interestingly, phylogenetic analysis using the D1/D2 region of LSU rRNA genes places three of the genera for which no –omics data is available (*Buwchfawromyces, Tahromyces,* and *Joblinomyces*) in a single distinct monophyletic clade (Figure 4). Future availability of –omics data is needed to confirm such topology. Finally, while the genus *Agriosomyces* has a distinct position in both ITS1 and LSU phylogenies (Figure 4, ITS), no clear association to any of the clades was apparent. As such, -omics data is hence needed to resolve the position of this genus.

### Rank assignment for supra-genus clades in the Neocallimastigomycota

Our analysis identifies and circumscribes four distinct clades in the *Neocallimastigomycota*. What taxonomic rank should be assigned to accommodate these clades? The Linnaean classification system places groups of genera into families. A recently proposed definition identifies fungal families as “a compilation of genera with at least one inherent morphological feature that they commonly share or which makes them distinct” [60]. The clades described in this study agree with such a definition, being a compilation of genera forming a distinct and monophyletic lineage with strong statistical support, and most of which share a common distinctive morphological feature (Table 2).

We propose retaining all currently described AGF genera in a single order (*Neocallimastigales*), and a single class (*Neocallimastigomycetes*) in the phylum *Neocallimastigomycota*. Such proposition is based on the lack of fundamental differences in their cellular structures, metabolic capabilities, ecological distribution, and life cycle phases in all currently described genera, coupled to the observed AAI values, when compared to the few studies utilizing this approach in fungi [51].

Beyond the four clades described above, we refrain from proposing an additional family for the D1/D2 LSU-defined and well-supported clade encompassing the genera *Buwchfawromyces, Tahromyces,* and *Joblinomyces* pending the availability of confirmatory phylogenomic data. As well, we refrain from proposing new families for the genera *Khyollomyces*, *Aklioshbomyces*, *Paucimyces*, and *Agriosomyces*, due to their current solitary positions in phylogenomic trees (Figure 1), although such proposition would be justified by the isolation of characterization of additional novel taxa and the availability of –omics data from such taxa. Such genera should be regarded as orphan taxa for the present time. The proposed novel families would be named after the first described genus within the clade: Clade 1 = *Neocallimastigaceae* comprising the genera *Neocallimastix* (Braune 1913 [61], Vavra and Joyon 1966 [62], Heath et al. 1983, [22]), *Ghazallomyces* (Hanafy et al. 2021) [5]*, Orpinomyces* (Breton et al. 1989 [63], Barr et al. 1989 [64]), *Pecoramyces* (Hanafy et al. 2017) [30], *Feramyces* (Hanfay et al. 2018 [31]), and *Aestipascuomyces* (Stabel et al. 2020, [34]); Clade 2 = *Caecomycetaceae* fam. nov., comprising the genera *Caecomyces* (Gold et al. 1988) [57] and *Cyllmayces* (Ozkose et al. 2001) [33], clade 3 = *Piromycetaceae* fam. nov., comprising the genus *Piromyces* (Gold et al. 1988) [57]; and clade 4 = *Anaeromycetaceae*, comprising the genera *Anaeromyces* (Breton et al. 1990) [65], *Capellomyces* (Hanafy et al. 2021) [5], *Liebetanzomyces* (Joshi et al. 2018) [66], and *Oontomyces* (Dagar et al. 2015) [29]. Such arrangement would necessitate amending the description of the family *Neocallimastigaceae*, currently encompassing all twenty genera, to include only the six genera stated above, rather than all twenty currently described AGF genera, as well as assigning the genera *Anaeromyces* (Breton et al. 1990), *Capellomyces* [5], *Liebetanzomyces* (Joshi et al. 2018) [66], and *Oontomyces* (Dagar et al. 2015) to the previously proposed (IF550425) nomenclature novelty family *Anaeromycetaceae*.

## Typification

### Emended description of fam. *Neocallimastigaceae*

Obligate anaerobic fungi with monocentric or polycentric thalli and filamentous rhizoidal system. Zoospores are polyflagellated in all described genera, with the exception of the monoflagellated genus *Pecoramyces*. The clade is defined by phylogenomic, phylogenetic and morphological characteristics. Currently accommodates the genera *Neocallimastix* (Braune 1913 [61], Vavra and Joyon 1966 [62], Heath et al. 1983, [22]), *Ghazallomyces* (Hanafy et al. 2021) [5]*, Orpinomyces* (Breton et al. 1989 [63], Barr et al. 1989 [64]), *Pecoramyces* (Hanafy et al. 2017) [30], *Feramyces* (Hanfay et al. 2018 [31]), and *Aestipascuomyces* (Stabel et al. 2020, [34]).

The emended description of the family *Neocallimastigaceae* is generally similar to that provided for the family *Neocallimastigaceae* [22], and order *Neocallimsatigales* [23], with amendments to exclude bulbous rhizoidal growth, and to circumscribe its boundaries to encompass a monophyletic clade of six genera. The clade is circumscribed by phylogenomic analysis, AAI values, and confirmed by LSU and RPB1 phylogenetic analyses, as well as morphological characteristics. The emended family encompasses the genera *Neocallimastix* (Braune 1913 [61], Vavra and Joyon 1966 [62], Heath et al. 1983) [22], *Orpinomyces* (Breton et al. 1989, Barr et al. 1989) [70, 71], *Pecramyces* (Hanafy et al 2017) [32], *Feramyces* (Hanafy et al 2018) [33], *Ghazallomyces* (Hanafy et al 2020) [5], and *Aestipascuomyces* (Stabel et al 2020) [8]. *Type genus*: *Neocallimastix* Braune 1913 [61], Vavra and Joyon 1966 [62], Heath et al. 1983, [22].

*Mycobank ID*: MB25486.

### Description of *Caecomycetaceae* fam. nov

Obligate anaerobic fungi that produce monoflagellated zoospores, monocentric or polycentric thalli that are either uni- or multisporangiate, and a bulbous rhizoidal system with spherical holdfasts. The clade is circumscribed by phylogenomic analysis, AAI values, and confirmed by LSU and RPB1 phylogenetic analyses, as well as morphological characteristics. Currently accommodates the genera *Caecomyces* (Gold et al. 1988) [57] and *Cyllmayces* (Ozkose et al. 2001) [33].

*Type genus: Caecomyces* (Gold et al 1988) [57].

*Mycobank ID*: MB844401

### Description of *Piromycetaceae* fam. nov

Obligate anaerobic fungi that produce monoflagellated zoospores, monocentric thalli, and filamentous rhizoidal system. The clade is circumscribed by phylogenomic analysis, AAI values, and confirmed by LSU and RPB1 phylogenetic analyses, as well as morphological characteristics. Currently accommodates the genus *Piromyces* (Gold et al. 1988) [57].

*Type genus*: *Piromyces* (Gold et al. 1988) [57].

*Mycobank ID:* MB844402

### Emended description of *Anaeromycetaceae* fam. nov

Obligate anaerobic fungi that produce monoflagellated zoospores, monocentric or polycentric thalli, and filamentous rhizoidal system. The clade is circumscribed by phylogenomic analysis, AAI values, and confirmed by LSU and RPB1 phylogenetic analyses, as well as morphological characteristics. Currently accommodates the genera *Anaeromyces* (Breton et al. 1990) [65], *Capellomyces* (Hanafy et al. 2021) [5], *Liebetanzomyces* (Joshi et al. 2018) [66], and *Oontomyces* (Dagar et al. 2015) [29].

*Type genus*: *Anaeromyces*, Breton et al. 1990 [65].

*Mycobank ID:* MB550425.

