## Supplementary document for "Phylogenomic analysis of the Neocallimastigomycota: Proposal of *Caecomycetaceae* fam. nov., *Piromycetaceae* fam. nov., and emended description of the families *Neocallimastigaceae and Anaeromycetaceae*"

| Table S1. Sequencing statistics | |  |  |  |  |  |  |
| --- | --- | --- | --- | --- | --- | --- | --- |
| **Genus** | **species** | **Strain** | **# of reads passing QC** | **# of transcripts** | **# of unique transcripts** | **# of predicted peptides** | **BUSCO (% completeness)** |
| *Aestapascuomyces* | *dupliciliberans* | R1 | 18050326 | 38274 | 35122 | 22092 | 97.35 |
| *Aklioshbomyces* | *papillarum* | WTS1 | 20122256 | 82655 | 76395 | 24167 | 96.52 |
| *Anaeromyces* | *contortus* | ABS23 | 17398956 | 30215 | 27650 | 16430 | 98.01 |
| *Caecomyces* | *communis* | FD27 | 16219646 | 106064 | 102476 | 18885 | 73.63 |
| *Caecomyces* | *communis* | TB33 | 16672181 | 115797 | 110482 | 26933 | 98.84 |
| *Capellomyces* | *forminis* | Cap2a | 15637321 | 38479 | 36152 | 17278 | 93.53 |
| *Cyllamyces* | *aberensis* | TSB2 | 20836243 | 30103 | 27264 | 17022 | 95.02 |
| *Khyollomyces* | *ramosus* | ZO44 | 21701226 | 37795 | 34026 | 17243 | 94.36 |
| *Liebetanzomyces* | *polymoprphus* | Orc37 | 20405825 | 29889 | 27171 | 18020 | 94.03 |
| *Neocallimastix* | *frontalis* | EC30 | 22468951 | 43578 | 39335 | 27248 | 94.36 |
| *Orpinomyces* | *joyonii* | AB3 | 18599007 | 24970 | 22972 | 16422 | 95.85 |
| *Orpinomyces* | *joyonii* | AB6 | 19777772 | 24180 | 22178 | 16132 | 97.84 |
| *Orpinomyces* | *joyonii* | ABC24 | 18199384 | 22649 | 20599 | 13858 | 97.68 |
| *Paucimyces* | *polynucleatus* | BB3 | 23272180 | 31463 | 28952 | 17911 | 98.18 |
| *Pecoramyces* | *ruminantium* | AS31 | 16958373 | 106687 | 103405 | 17117 | 81.26 |
| *Pecoramyces* | *ruminantium* | AS32 | 20268027 | 23347 | 21844 | 15996 | 89.22 |
| *Pecoramyces* | *ruminantium* | Orc32 | 23381792 | 35580 | 31975 | 23021 | 99.00 |
| *Piromyces* | *cryptodigmaticus* | Axs23 | 21276025 | 45838 | 41760 | 21709 | 92.21 |
| *Piromyces* | *finnis* | DonB11 | 21413984 | 26131 | 23413 | 17455 | 99.50 |
| *Piromyces* | sp. NZB19 | Ors32 | 23819290 | 43652 | 39201 | 28405 | 99.50 |

**Figure S1.** Upper triangle matrix (A) and box and whisker plots (B) for the alignment divergence values obtained for all possible pairs of sequences in Figure 3. (A) Isolate names in rows and columns are color coded by clade (clade 1, purple; clade 2, lavender; clade 3, orange; clade 4, light blue). The alignment divergence values for each clade are shown within a thick border. Intra-genus values are shown in red text with pink highlight, intra-clade/ inter-genus values are shown in blue text with light blue highlight, while inter-clade values are shown in green text with light green highlight. *Ghazallomyces constrictus* was included in clade 1 and *Oontomyces anksri* was included in clade 4 as explained in the main text. Values for the genera unaffiliated with the 4 clades are highlighted in grey. (B) Box and whisker plots constructed using the values in (A). Intra-genus values (red) are shown both including and excluding the genus *Piromyces*. Intra-clade/ inter-genus values are shown in blue. Inter-clade values are shown in green. Each box plot spans the region between the 25-percentile to 75-percentile, while the whiskers limit the minimum and maximum scores excluding the outliers. The thick line inside the box marks the median, while the ‘x’ corresponds to the average value.

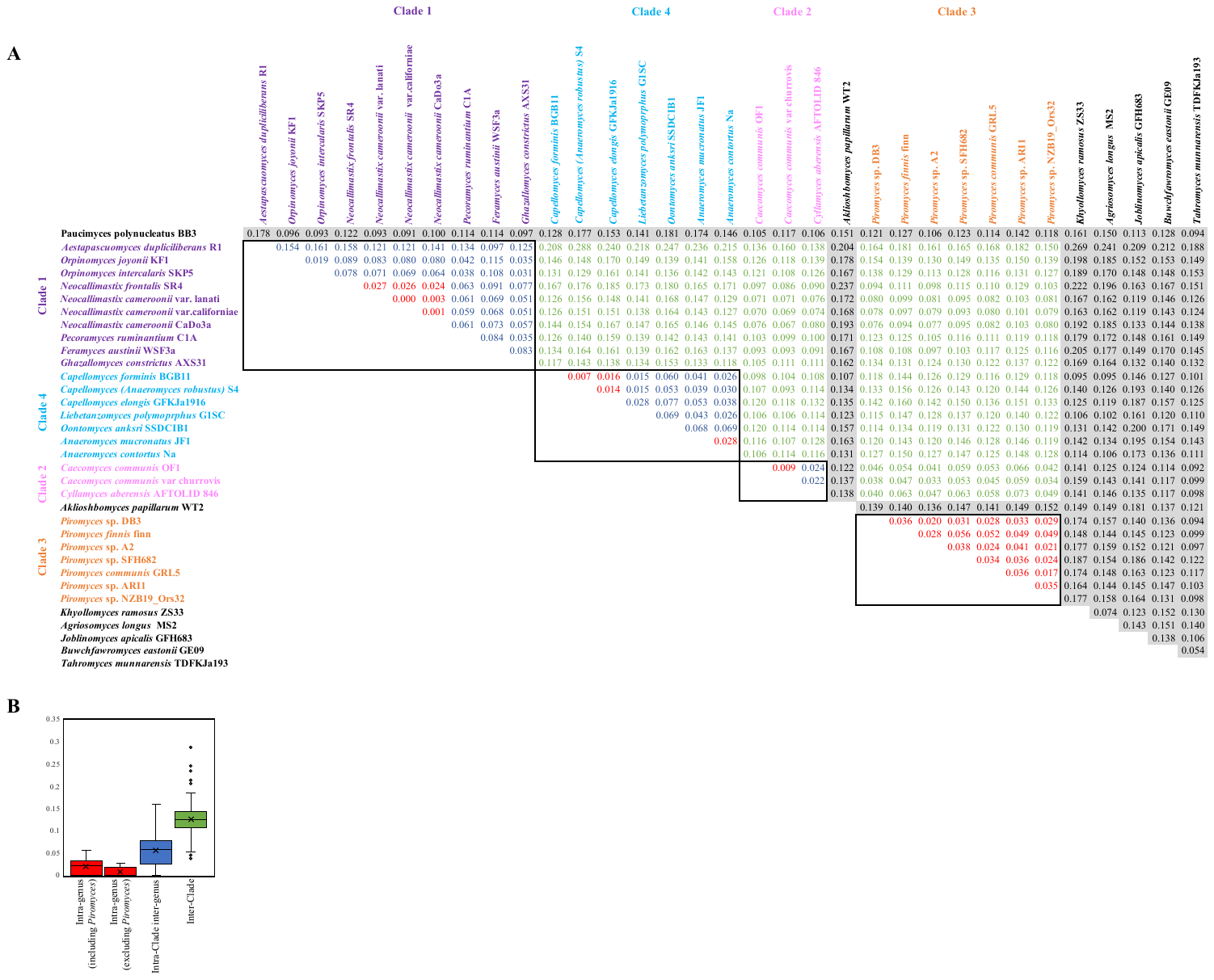

**Figure S2.** Upper triangle matrix (A) and box and whisker plots (B) for the alignment divergence values obtained for all possible pairs of sequences in Figure 5. (A) Isolate names in rows and columns are color coded by clade (clade 1, purple; clade 2, lavender; clade 3, orange; clade 4, light blue). The alignment divergence values for each clade are shown within a thick border. Intra-genus values are shown in red text with pink highlight, intra-clade/ inter-genus values are shown in blue text with light blue highlight, while inter-clade values are shown in green text with light green highlight. Values for the genera unaffiliated with the 4 clades are highlighted in grey. (B) Box and whisker plots constructed using the values in (A). Intra-genus values (red) are shown both including and excluding the genus *Piromyces*. Intra-clade/ inter-genus values are shown in blue. Inter-clade values are shown in green. Each box plot spans the region between the 25-percentile to 75-percentile, while the whiskers limit the minimum and maximum scores excluding the outliers. The thick line inside the box marks the median, while the ‘x’ corresponds to the average value.

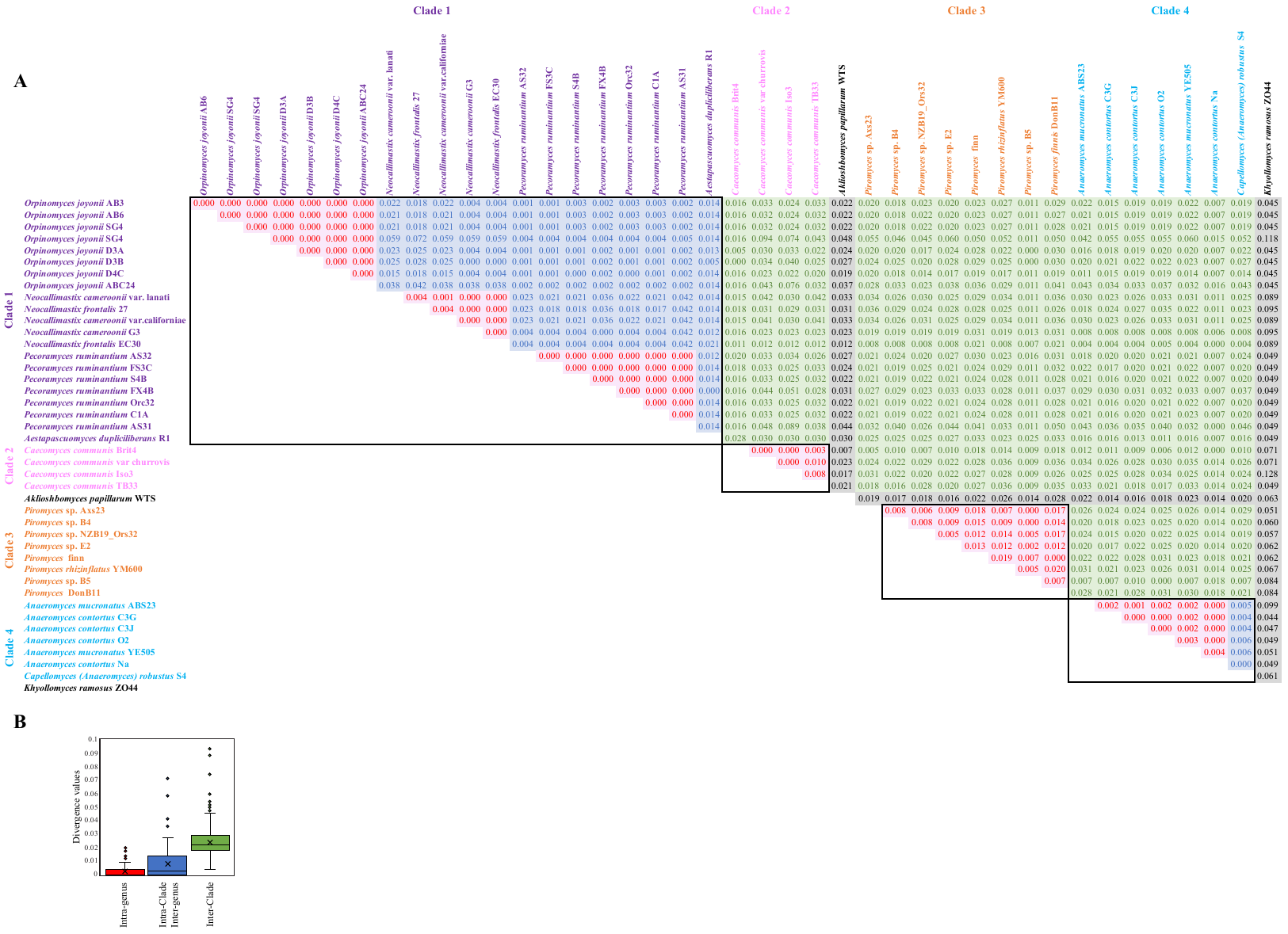
